# Physical biology of cell-substrate interactions under cyclic stretch

**DOI:** 10.1101/2020.10.23.350959

**Authors:** Siddhartha Jaddivada, Namrata Gundiah

**Author notes:** For correspondence: Namrata Gundiah, Biomechanics Laboratory, Department of Mechanical Engineering, Indian Institute of Science, Bangalore 560012. **Email**, **Tel (Off/ Lab)**: 91 80 2293 2860/ 3366.

## Abstract

Mechanosensitive focal adhesion complexes mediate the dynamic interactions between cells and substrates, and regulate cellular function. Integrins in adhesion complexes link substrate ligands to stress fibers in the cytoskeleton, and aid in load transfer and traction generation during cell adhesion and migration. A repertoire of signaling molecules, including calcium, facilitate this process. We develop a novel one-dimensional, multi-scale, stochastic finite element model of a fibroblast on a substrate which includes calcium signaling, stress fiber remodeling, and focal adhesion dynamics that describes the formation and clustering of integrins to substrate ligands. We link the stochastic dynamics involving motor-clutches at focal adhesions to continuum level stress fiber contractility at various locations along the cell length. The stochastic module links to a calcium signaling module, *via* IP_3_ generation, and adaptor protein dyanamics through feedback. We use the model to quantify changes in cellular responses with substrate stiffness, ligand density, and cyclic stretch. Results show that tractions and integrin recruitments vary along the cell length and depend critically on interactions between the stress fiber and reversibly engaging adaptor proteins. Maximum tractions and integrin recruitments were present at the lamellar regions. Cytosolic calcium increased with substrate stiffness and ligand density. The optimal substrate stiffness, based on maximum tractions exerted by the cell, shifted towards stiffer substrates at high ligand densities. Cyclic stretch increased the cytosolic calcium and tractions at lamellipodial and intermediate cell regions. Tractions and integrin recruitments showed biphasic responses with substrate stiffness that increased with ligand density under stretch. The optimal substrate stiffness under stretch shifted towards compliant substrates at a given ligand density. Cells deadhere under stretch, characterized by near-zero recruitments and tractions, beyond a critical substrate stiffness. The coupling of stress fiber contractility to adhesion dynamics is essential in determining cellular responses under external mechanical perturbations.

**Statement of Significance:** Cells are exquisitely sensitive to substrate ligand density, stiffness, and cyclic stretch. How do cell-substrate interactions change under cyclic stretch? We use a systems biology approach to develop a one-dimensional, multi-scale, stochastic finite element model of cellular adhesions to substrates which includes focal adhesion attachment dynamics, stress fiber activation, and calcium signaling. We quantify tractions along the cell length in response to variations in substrate stiffness, cyclic stretching, and differential ligand densities. Calcium signaling changes the stress fiber contractility and focal adhesion dynamics under stretch and substrate stiffness. Cell tractions and adhesions show a biphasic response with substrate stiffness that increased with higher ligand density and cyclic stretch. Chemomechanical coupling is essential in quantifying mechanosensing responses underlying cell-substrate interactions.

## Introduction

Cyclically loaded cells, such as arterial fibroblasts, endothelial cells, and smooth muscle cells, must constantly adapt to dynamically changing mechanical loads. Cell remodeling under stretch is vital to the activation of complex biochemical signaling pathways that orchestrate cellular functions related to contractility, differentiation, growth, and migration over a short duration (1–3). Mechanotransduction over a longer duration regulates gene expression and protein synthesis that may, over time, cause changes to the underlying extracellular matrix (ECM) properties and contribute to the progression of diseases such as fibrosis and aneurysm growth (4, 5).

Heterodimeric transmembrane proteins, integrins, are an important component of focal adhesion (FA) complexes and mediate the complex interactions between cells and substrates (4). Integrins attach to the ECM through their extracellular domains, connect to cytoskeletal stress fibers (SF) *via* intracellular domains, and permit bi-directional signaling between the cell and the substrate (6). Several distinct, multi-protein assemblies in the FA complexes contribute to the downstream cell signaling cascades (7–9). Scaffolding and adaptor FA proteins, such as vinculin, paxillin, talin, and zyxin, link the SF to the ECM *via* integrins, whereas signaling proteins are locally recruited to generate and mediate the development and maturation of FA under mechanical stimuli (7, 10, 11).

Integrins cluster and corral into adhesomes (12) in the presence of extracellular magnesium from the ECM (13) and cytoplasmic forces (14). Forces cause a conformational change in the integrin structure, altering them from a *bent* state to an *extended* one, and initiate downstream signaling when coupled to the SF’s (15). Reinforcement and maturation of integrins link them to proteins, such as talin, tensin, α-actinin, and vinculin, collectively called *adaptor proteins,* which increase the size, shape, and molecular composition of the adhesome (16, 17). These changes are in response to forces applied by the SF or through cyclic substrate stretch (18). Fibroblasts under uniaxial cyclic stretch reorient in a direction orthogonal to the stretch direction at high frequencies (>1 Hz) and amplitudes over 5% (19–21). Tendon fibroblasts express higher levels of α-smooth muscle actin and phospholipase A2 when oriented parallel to the stretching direction (20). Cell responses during stretch require a change in the integrin density (22). FRAP experiments show actin incorporation at the FA’s and along the entire length of the SF (23). Growth and remodeling of SF’s are hypothesized to be essential to the re-orientational responses of cells under cyclic stretch (24).

Chan and Odde modeled the physical coupling of cells on deformable substrates using spring-like molecular clutches, representing adaptor proteins that generate tractions and stochastically form or break (25–26). Individual clutches reversibly engage to the substrate resulting in tension build-up in successfully engaged clutches that link to the cytoskeletal actin. Myosin motors cause a retrograde flow of actin and work against the resistance provided by the engaged clutches. This interaction results in two modes of interaction: a stalled mode that generates high tractions and adhesions, and a load-and-fail mode with lower tractions and adhesions (26). The model demonstrated an optimum substrate stiffness based on the maximum tractions exerted by cells on substrates. The model did not however incorporate chemical signaling, associated with integrins, and the corresponding changes in SF activation caused by clutch engagements.

Cell traction measurements on substrates show an initial monotonic increase with increasing substrate stiffness and higher ligand affinity; the FA size also increased linearly with force (15, 27). Deshpande and colleagues developed a generalized model to represent the SF crossbridge dynamics and quantified the cellular response to varied substrate compliance (28). Strong spatial gradients in SF contractility are present along the cell length through coupled Rho signaling at integrin attachment sites (29). Cyclic stretching of dermal fibroblasts causes upregulation in the expression levels of F-actin capping protein a-1 and myosin; this suggests increased cellular tractions in cyclically stretched cells (30). Higher tractions in cyclically stretched cells is accompanied with increased cell stiffness (24) and intracellular calcium transients *via* stretch – activated channels (31, 32). Increased calcium flux under stretch also depends on the activation of phosphatidyl-inositol pathway, induced by membrane perturbations, which suggests a possible role for FA clustering (31).

How do dynamic changes in integrin recruitments, coupled to SF contractility, result in the development of tractions in cyclically stretched cells? We develop a novel 1D multi-scale model of a fibroblast to include biochemical signaling and remodeling of the cytoskeleton and FA dynamics at different locations along the cell length. We quantify changes to the SF contractility, FA remodeling, and chemical signaling in response to external cyclic mechanical stretch. We also test how cellular responses are altered with substrate stiffness and ligand density which is relevant in the context of tissue fibrosis. Our results show the importance of calcium dynamics in delineating differences in the cell tractions under cyclic stretch. Chemomechanical coupling of the SF and FA dynamics is also useful to characterize the individual roles of adaptor proteins, such as talin, in mechanosensing.

## Methods

### A computational model of cell contractility and adhesion formation

We used a systems biology approach to model cell-substrate interactions under cyclic stretch. Deformations during cell-substrate interactions were calculated using a one-dimensional stochastic finite element method (SFEM). The substrate and SF were discretized into 14 and 28 elements respectively (Fig. 1A) to explore spatial differences in FA and tractions under static and stretch conditions. The cell includes a lamellipodia region, located between nodes 1-3 and 27-29, and lamellar regions, located at nodes 3-9 and 21-27, on either side of the cell center. Ligands with a uniform concentration on the substrate attach to integrins in the cell membrane and connect the other structural adaptor proteins (clutches) to the contractile SF (Fig. 1B). The model consists of different modules that couple the mechanics at hierarchical scales ranging from the stochastic dynamics of individual clutches to the continuum level SF contractility that includes calcium signaling.

**Figure 1.**
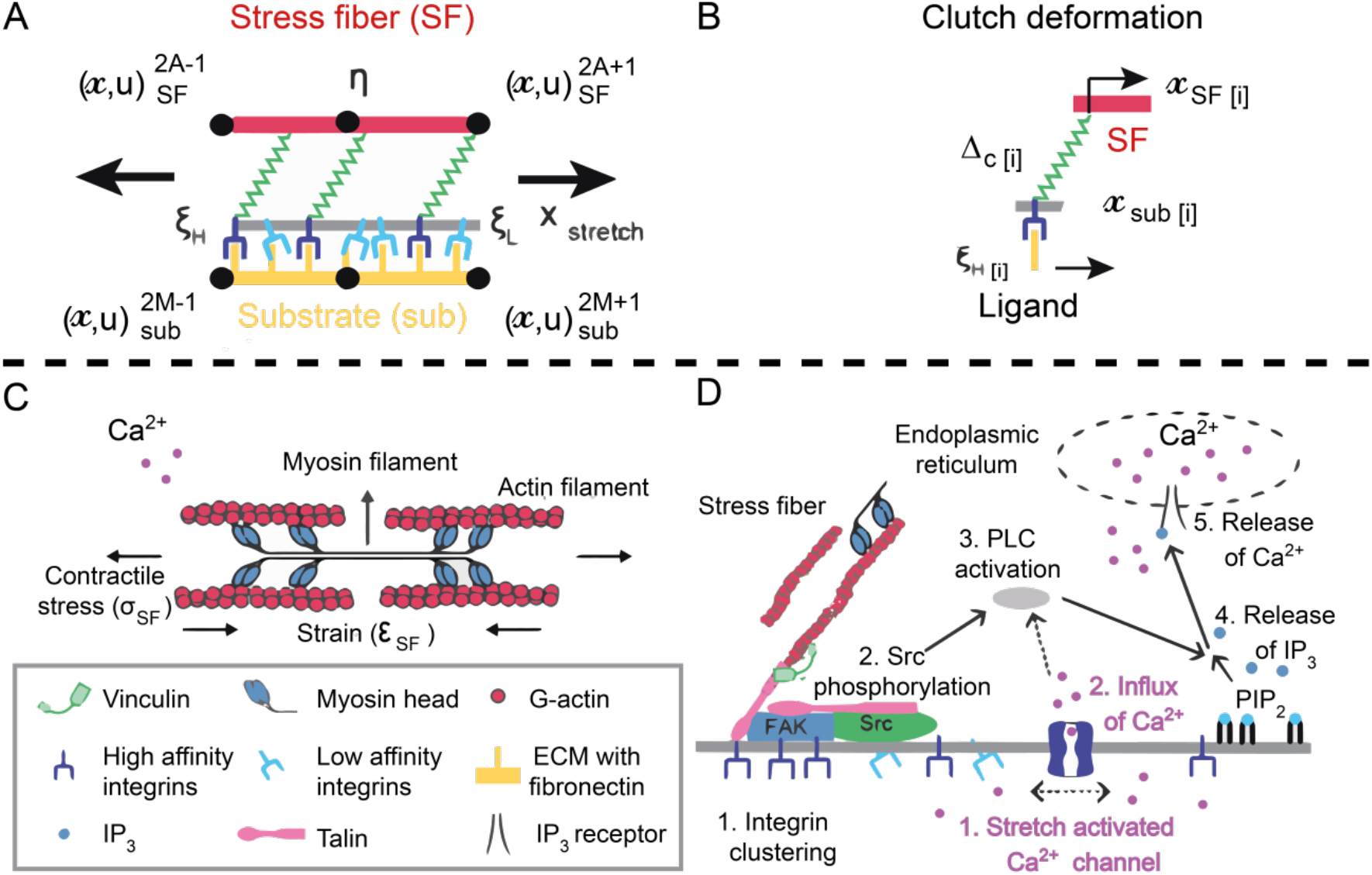
**(A)** A representative FE element between SF nodes 2A-1 and 2A+1 and substrate nodes 2M-1 and 2M+1 is shown. The cell membrane (gray) with integrins attaches to a ligand coated substrate (yellow). 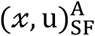 and 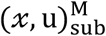 are nodal positions and displacements for the SF and substrate respectively. (**B**) Adaptor proteins (green) attach the i^th^ ligand on the substrate through high-affinity integrin attachments to the SF’s (red) with activation, η^A^. Deformations, Δ_*c*[*i*]_, between the clutch attachment points at the SF,*x*_*SF*[*i*]_, and the substrate, *x*_*sub*[*i*]_, results in changes in concentrations of high-affinity, *ξ*_*H*_, and low-affinity integrins, *ξ_L_*, at the cell membrane. The concentration of high-affinity integrins at the bottom end of the clutch is represented as *ξ*_*H*[*i*]_. **(C)** Actomyosin interactions, through cross-bridge cycling, induce contractile stress (***σ***_*SH*_) and strain (***ε_SF_***) in the presence of calcium. (**D**) Calcium signaling in the model has two possible pathways that include integrin clustering (black) or result due to membrane stretch (purple). Both pathways result in PLC activation in the cell.

The stochastic module includes the engagement/ disengagement of clutches between the SF and the substrate characterized with Young’s modulus, Esub, and containing a uniform fibronectin coating with defined ligand densities, nc (25–27). Cell-substrate interactions are delineated into four different phases: (i) formation of reversible bonds between high-affinity integrins and ligands, (ii) clustering of high-affinity integrins at FA complexes, (iii) recruitment of structural adaptor proteins by integrin clusters in the FA, and (iv) contractility due to SF attachment to adaptor proteins (33, 34). SF contractility due to cross-bridge cycling between actin and myosin requires calcium (Fig. 1C) and results in deformations of adaptor proteins and the ligand coated substrate. Both integrin dynamics at FAs and stretch-dependent ion channels are associated with calcium flux in the cell (Fig. 1D) that is described using a signaling module (31, 35). Calcium feedback is especially important in the context of cells under cyclic stretch (31, 32). We will next describe the biochemical and mechanical coupling between these different components in the model (Fig. 2).

**Figure 2:**
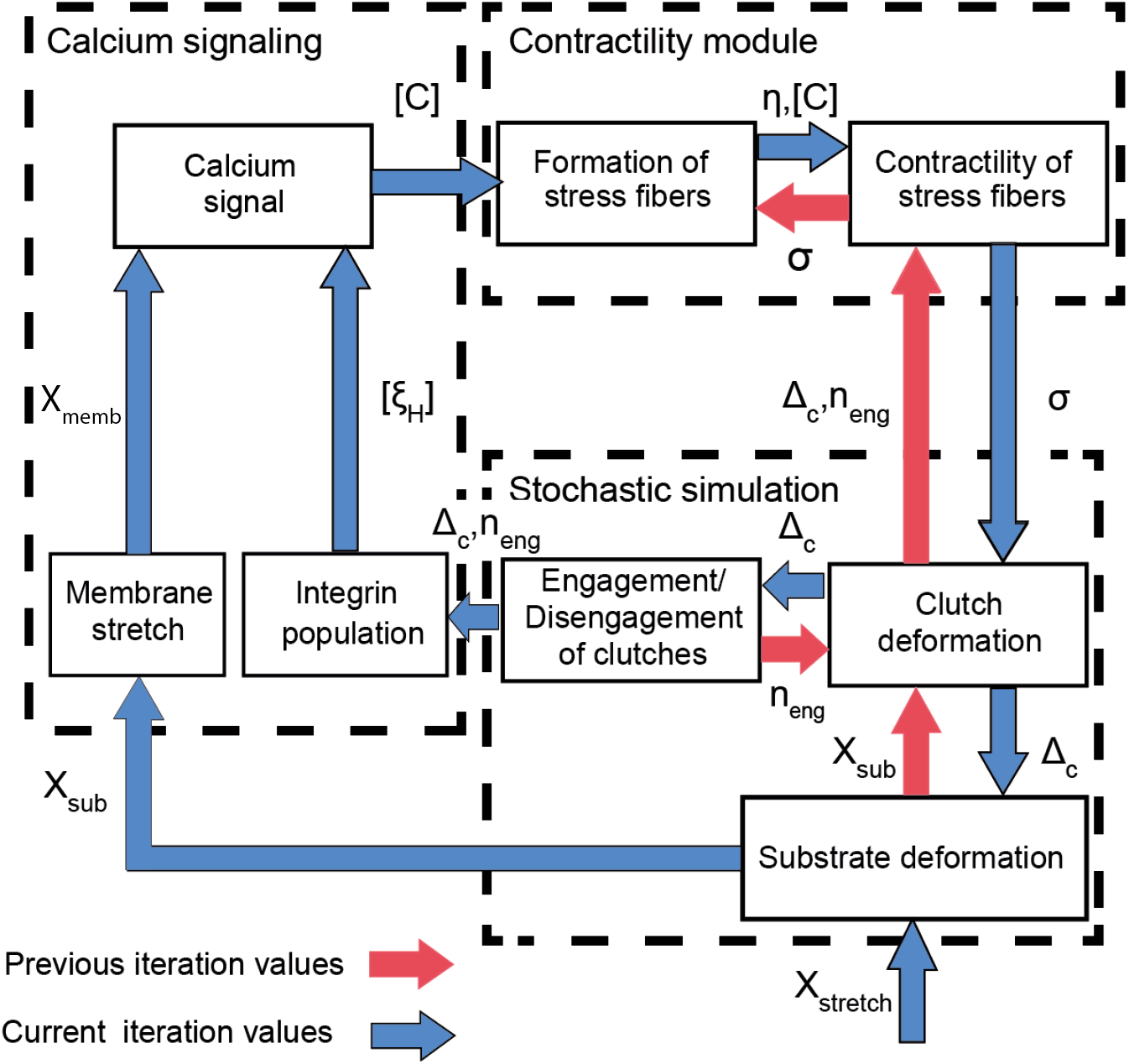
The cell-substrate interaction model includes stochastic simulations, SF contractility, and calcium signaling modules. Interactions between the different modules are indicated by arrows for the previous and current iteration along with the corresponding variables. Clutches in the cell membrane reversibly engage (n_eng_) or disengage with the substrate and are modeled through the stochastic module. SF activation (η) in the presence of calcium results in stress (*σ_SF_*) that causes deformation (Δ_c_) of the engaged clutches, and a corresponding substrate displacement (*u_sub_*) that is simulated using the contractility module. *u_sub_* and Δ_c_ causes activation of calcium signal through membrane stretch and aggregation of high-affinity integrins (ξ_H_) obtained using the calcium signaling module. The change in calcium (*C*) is an input in the contractility module.

#### 1. Motor-clutch stochastic dynamics

We use the motor-clutch model to simulate interactions between the SF and ligands attached uniformly to a linearly elastic ECM substrate (Fig. 1A). Ligands on the substrate in each element are in a disengaged state at the start of the simulation; these engage/ disengage stochastically to the SF through clutches (26). Fig. 1A shows a representative SF element with clutches that reversibly bind with rate, *k*_*on*_[*i*]__, to the SF and the i^th^ ligand on the substrate and disengage with a force-dependent rate, *k*_*off*_[*i*]__ given by the Bell model for slip bonds (36) as

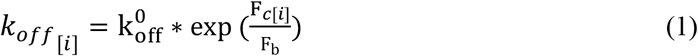

*F_b_* is the characteristic rupture force, and F_*c*[*i*]_ is the force transmitted by the *i^th^* clutch to the substrate in element A located on the SF, given by

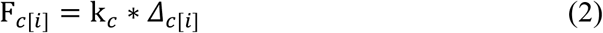

where k_c_ is the stiffness of the clutch in this expression.

Clutches resist the actin retrograde flow caused by active stress, *σ_SF_*, generated in the SF. Engaged clutches are subject to deformations, *Δ*_*c*[*i*]_, due to the active SF stress. The dynamics of clutch engagements/ disengagements were obtained using Monte Carlo simulations. In this method, the ensemble of clutches at each element of the cell were initially assigned uniform random numbers (0 ≤ *URN*_[*i*]_ ≤ 1). The off-rate for each engaged clutch was determined using deformations in Eq. 1. The event times for engagement, *t*_*on* [*i*]_, or disengagement, *t*_*off*[*i*]_, of the ligand, i, in element A were calculated using

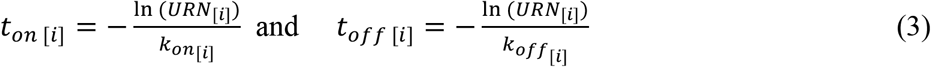

The position *x*_*SF*[*i*]_ of engaged clutches 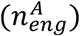 in the SF element from the previous iteration was updated (25) over the minimum event time (*t_on/off_* = min (*t*_*on* [*i*]_/*t*_*off* [*i*]_)).

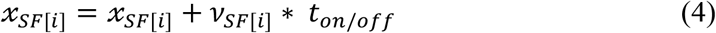

*v*_*SF* [*i*]_ is the rate of deformation of the SF at the clutch attachment point which was calculated using the FEM. The clutch deformation, *Δ*_*c* [*i*]_, was determined using the difference in coordinates of the clutch attachment points on the SF, *x*_*SF*[*i*]_, and the substrate, *x*_*sub*[*i*]_. Clutch deformation is also equivalent to the difference in the displacements of attachments at SF, u_*SF*[*i*]_, and the substrate, U_*sub*[*i*]_

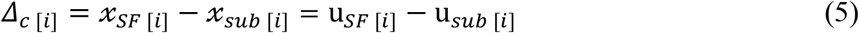

The transmitted force, *F*_*c*[*i*]_, and cyclic stretch, *x_stretch_*, results in a local substrate displacement, 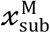. The corresponding change in the position of ligand attachments between nodes 2M-1 and 2M+1 of the substrate, *x_sub_[*i*]* was determined using interpolation. Mechanical stiffness of the clutches, *k_c_*, and the substrate, *E_sub_*, determine the resistance to clutch loading by the SF.

#### 2. Contractility module

A continuum-level contractility module was used to quantify the SF activation kinetics and is characterized through the coupling of the calcium-dependent cross-bridge cycling (Fig. 1C) and the tension-dependent SF assembly/ disassembly. We use a first-order rate equation to parametrize the assembly/disassembly of SF bundles using a non-dimensional activation parameter, η, at each location in the cell given by (36)

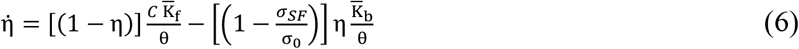

*η* is the ratio of the concentration of polymerized actin and phosphorylated myosin in the SF to their maximum possible concentrations. 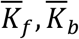 are the rate constants that govern the formation and dissociation of SF’s with a time constant, θ. *C* is the concentration of calcium, *σ_SF_*, is the stress in the SF and *σ*_0_ is the isometric stress of the SF. Eq. 6 shows that the rate of SF assembly decreases with SF activation at each element, A, and is proportional to calcium availability. In contrast, SF dissociation is directly proportional to the activation at each node and is a function of the SF stress. The dissociation rate is zero when the fibers are held at their isometric stress, *σ*_0_, at each element and increases linearly at lower stress.

The choice of isometric stress value is an important aspect of the simulation. We assume that the isometric stress of the SF at each element, *σ*_0_, is directly proportional to the activation level of the SF, σ_0_ = η*σ_max_* where, *σ_max_* is the isometric stress during the maximum possible concentration of the polymerized actin and phosphorylated myosin in the SF. The value of *σ_max_* was taken to be a product of the myosin motor concentration, *n_m_* and maximum force generated by individual motors, *F_m_*. *σ_max_* in each element is generated by cross-bridge cycling similar to that in muscle cells (Fig. 1C). We used a Hill-like relation to model the influence of tension in the SF based on the rate of extension/shortening of SF given by

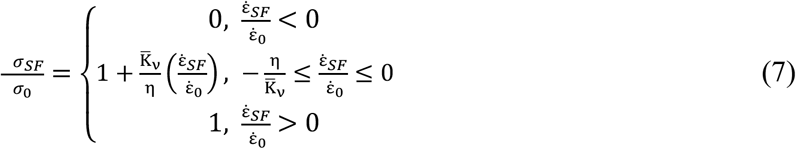

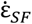 is the rate of change in the SF length,which is positive for lengthening and negative for shortening. The non-dimensional constant, 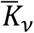, is the fractional reduction in stress when the shortening rate increases by the isometric value, 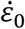.

#### 3. Biochemical coupling of calcium signaling

Cyclic stretching of the membrane causes the release of calcium ions that alters the SF contractility (35). Calcium flux in the cell is a consequence of two signaling pathways: first, through changes in the concentration of high-affinity integrins, *ξ_H_*, at the base of engaged clutches. Second, membrane stretch, *u_memb_*, causes activation of stretch-activated calcium ion channels that results in downstream signaling to alter the calcium concentration. (Fig. 1D). Membrane stretch is assumed to be proportional to the displacement, *u_sub_*, in the substrate.

We calculate the density of high-affinity integrins, at ligand i in SF element A, every iteration of the Monte-Carlo simulation, as a ratio of the reference concentration of integrins, *ξ_R_*, to the change in chemical energy due to integrin conformation change (37)

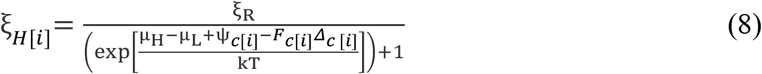

The denominator includes terms for the difference in chemical potentials between high and low-affinity integrins (*μ_H_* — *μ_L_*), stretch energy in the integrin – ligand complex for each ligand (ψ_*c* [*i*]_), and the work conjugate (*F*_*c* [*i*]_*Δ*_*c*[*i*]_) term. Eq. 8 is used to account for integrin clustering that is altered in the presence of variable clutch force induced by SF and substrate stretch.

High-affinity integrins initiate the production of signaling molecule, IP_3_, that diffuse and get de-phosphorylated within the cytosol. IP_3_ molecules induce the opening of calcium gated channels in the endoplasmic reticulum (ER) and cycle back into the ER through molecular pumps. The rate change in the concentration of IP_3_, S, at a specific location, *x_i_*, due to diffusion is given using a reaction-diffusion equation by (37)

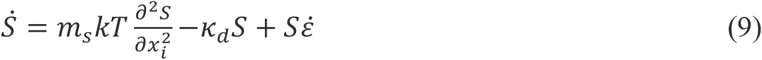

*m_s_* represents the motility of IP_3_ in the cell, k is the Boltzmann’s constant, and T is the absolute temperature. The rate equation also includes the de-phosphorylation of IP_3_ described with a reaction rate, *k_d_*. The last term in the equation is used to account for the effects of cell stretch on calcium dynamics *via* the strain rate, 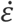. The flux boundary condition is given by

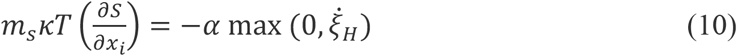

An increase in *ξ_H_* with time thus results in the production of IP_3_ with a non-dimensional proportionality constant α (37).

The concentration of calcium ions in the cytosol is normalized with respect to the maximum calcium in cells to *C* (0≤*C*≤1). The rate equation for *C* is given by assuming first-order kinetics as,

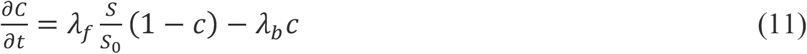

where *λ_f_* is the rate constant governing the rate of release of calcium ions from the IP_3_-gated reserve in the endoplasmic reticulum, and *S*_0_ is a reference concentration of IP_3_. Calcium ions increase with increasing S, and their reabsorption into the ER is governed by another rate constant, *λ_b_*, and the calcium concentration, *C*. These equations are useful to model the role of integrin populations and membrane stretch on the overall calcium that can vary at each point within the cell.

#### 4. Coupling between the chemomechanical modules to simulate cell-substrate interactions during static and stretch conditions

We simulate the cell-substrate interactions through the SFEM method implemented in MATLAB (v8.2 2013a; The Math Works, Natick, MA). The simulation involves temporal integration of three coupled modules through a staggered approach and a subsequent implementation of mechanical equilibrium in the SF, clutches, and the substrate at the end of each time step. The modules embody stochastic clutch dynamics, calcium signaling, and SF contractility during cell-substrate interaction. Feedback between the different modules involving current and earlier iteration values are indicated in Fig. 2. Mechanical equilibrium of the cell-substrate system is implemented through the following set of coupled equations

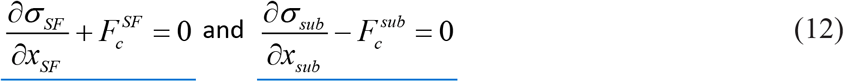

*x_SF_* and *x_sub_* are the coordinates along the length of cell and substrate domains, respectively. *σ_SF_* is the stress in SF specified by constitutive Eqs. 6 and 7, and *σ_sub_* is the stress in the hyperelastic substrate specified as:

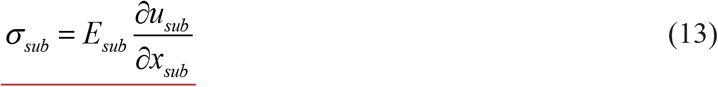

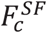 and 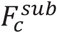 are the forces transferred by clutches to SF and substrate respectively. These forces are the summation of clutch forces F_*c*[*i*]_ calculated using Eqs 2 and 5:

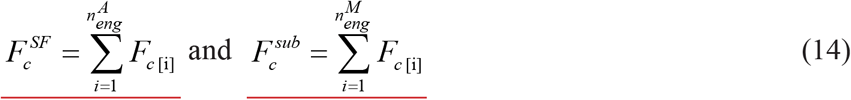

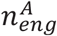 and 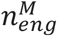 are the number of clutches engaged to SF and substrate respectively at a given location.

Deformations in the cell and substrate were obtained by establishing a weak form of Eq. 12 using the Galerkin procedure. Boundary conditions for SF are given by *σ_SF_* = 0 and *u_SF_* = 0 at *x_SF_* = ± *L_c_* and *x_SF_* = 0, respectively. The boundary condition for the substrate is given by *u_sub_* = 0 at *x_sub_* = 0 and *u_sub_* = 0 at *x_sub_* = ± L_s_/2, and for substrate under stretch *u_sub_* = ± (0.1 x *L_s_* /4) x (1+ sin(2π t- π/2)) at *x_sub_* = ± L_s_/2.

The variational form of Eq 12 is given as

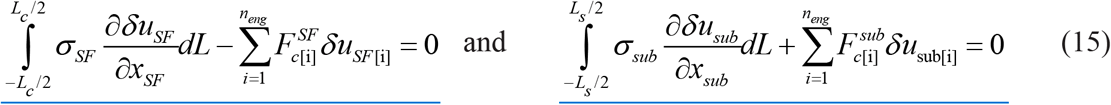

*n_eng_* is the total number of clutches engaged.

The linearized form of Eq 15 at time t+*Δ*t is given by

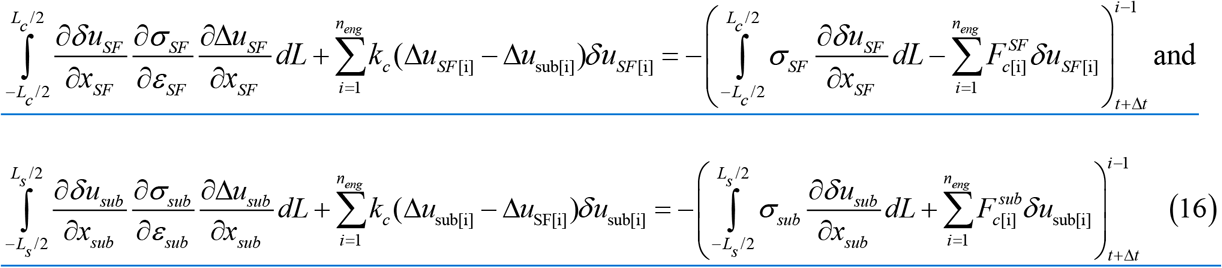

The variational equations in Eq 16 were discretized through 1D quadratic on cell and substrate domains (*H_SF_* and *H_sub_*) respectively. The displacements in domains were interpolated as 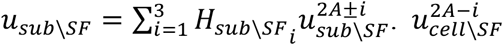 are the displacements at the nodes as represented in Fig 1. Strains in each SF element (*ε^A^*) and substrate element (*ε^M^*) were evaluated using

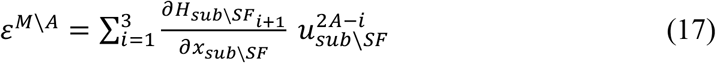

Substitution of interpolations in Eq 16 yielded the following discretized equation:

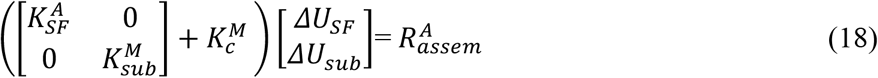

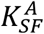 and 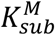 are the global assembly element stiffness matrices 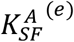 and 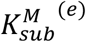 of SF and substrate, respectively

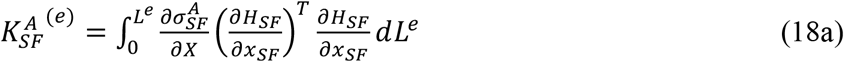

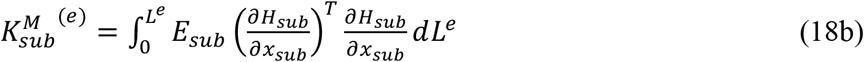

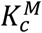 is the stiffness contribution of *n_eng_* clutches engaged between SF elements and substrate elements obtained by global assembly of 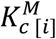. This term represents the contribution of the i^th^ clutch attached at local coordinates 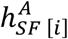 and 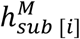. These contributions were assembled into 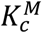, based on corresponding indices of SF (A) and substrate (M) elements.

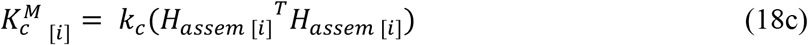

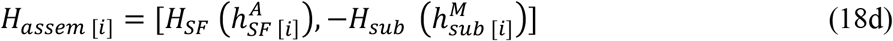

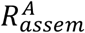 is the internal residual force of the cell-substrate interaction which constitutes clutch forces 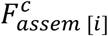 and, the elemental internal force in SF and substrate, 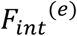. Both 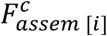 and 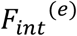 were assembled into global matrices, 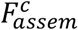 and *F_int_*, based on the associated element indices.

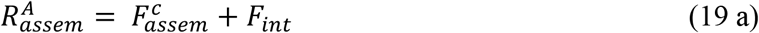

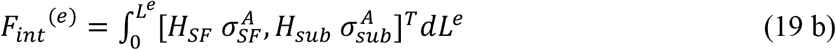

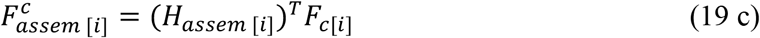

The increments in displacements at nodes of the cell and substrate *ΔU_sub/SF_* were calculated from Eq. 18 using the Newton-Raphson method. The coordinates of attachment of clutches to the SF, *x*_SE[*i*]_, were updated using Eq. 4.

The SF activation, η, was evaluated at every time step, Δt, to determine the SF contractile stress, *σ_SF_*, using Eqs 6 and 7 in the contractility module. SF contractility results in clutch deformations, *Δ*_*c*[*i*]_. SF and substrate displacements, *u_SF/sub_*, were updated using deformation increments, *ΔU_SF/sub_*, calculated using Eq 18. Forces in the clutches, *F*_c[*i*]_, were determined using Eqs 2 and 5. The clutch deformations were used to determine the engagement/disengagement rates of clutches in the stochasticity module using Eqns. 1 and 3. The clutch deformation Δ_c[i]_, clutch forces *F*_c[i]_, and membrane deformation, *u_memb_* = *u_sub_*, were input in the calcium signaling module. The population density of high affinity integrins at the end of engaged clutches, ξ_H[i]_, was determined from Eq. 8 using both *Δ*_c[i]_ and *F*_c[i]_. The rate of change in high-affinity integrins was calculated to determine IP_3_ production (Eq. 9). The discretized versions of both Eqs. 9 and 11 were integrated using 8^th^ order Runge-Kutta method to update IP_3_ concentration (5) and calcium level *(C)*. Eq. 1–11 and 19 were temporally integrated until the simulation time *t_end_* is reached. A flow chart for the implementation of the algorithm is included in Fig S1.

#### 5. Parameters used in the simulation

Table 1 summarizes the simulation parameters for each of the different modules and the corresponding sources from which these values were obtained. The length of the fibroblast cell, L_C_, is assumed to be 60 μm. The reference strain rate 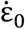 is selected to be 4 x 10^-3^ so that retrograde velocity of 120 nm/s is obtained at the cell edge. The values of myosin concentration (n_m_) and maximum SF stress (σ_max_) were selected such that the magnitude of tractions match experimental data for fibroblasts (10, 15). The velocity reduction constant value, 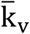, was selected such that the velocity relation used in Chan and Odde (25) was reproduced in the Hill’s equation (Eq. 7). Substrate stiffness, k_sub_, was in the range 10-1000 kPa to represent a range of tissue stiffness observed *in vivo.* The rate constant for SF formation (k_f_) was chosen to be higher than the dissociation constant (k_b_) such that SF activation was possible in the absence of stress.

**Table 1:**
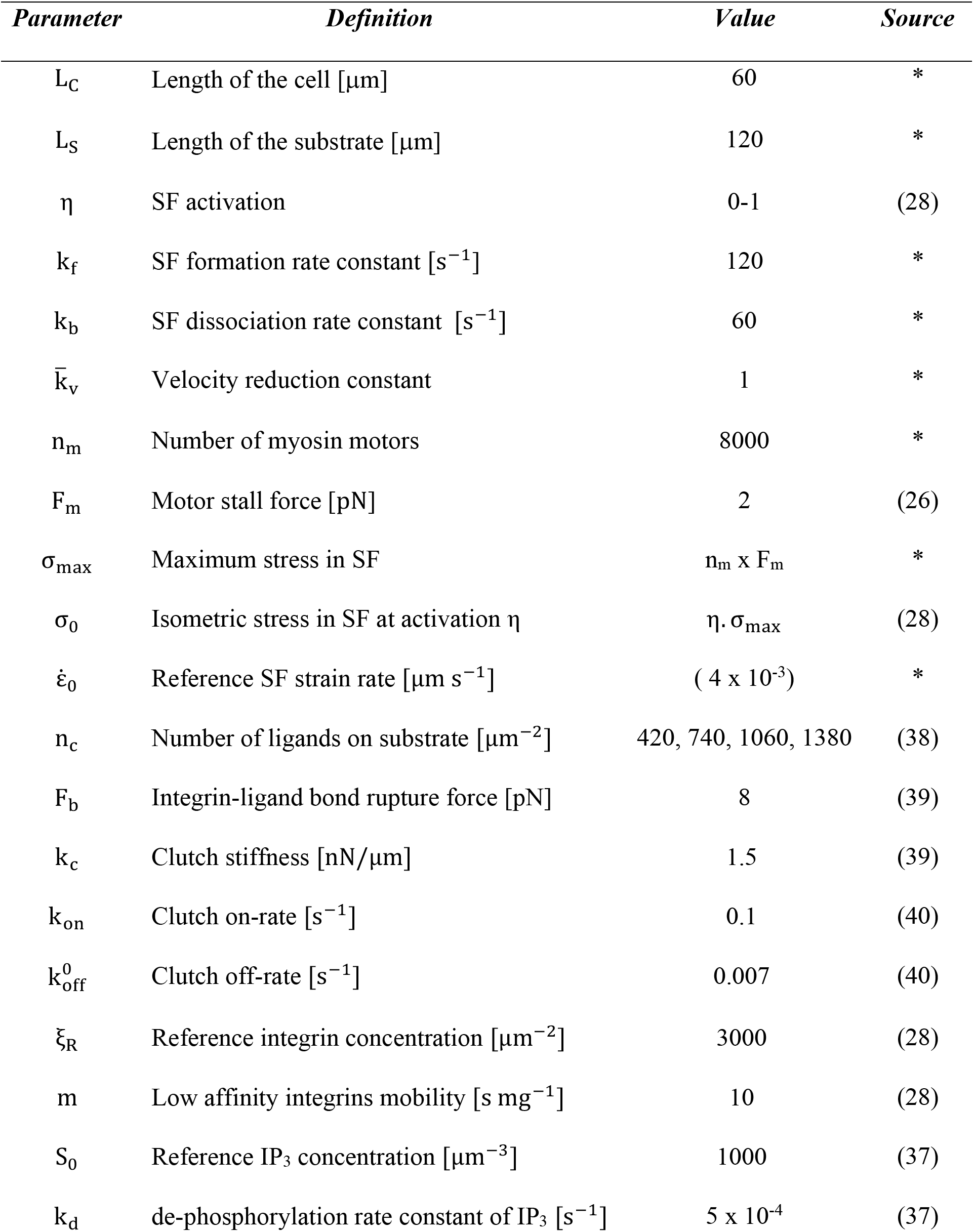
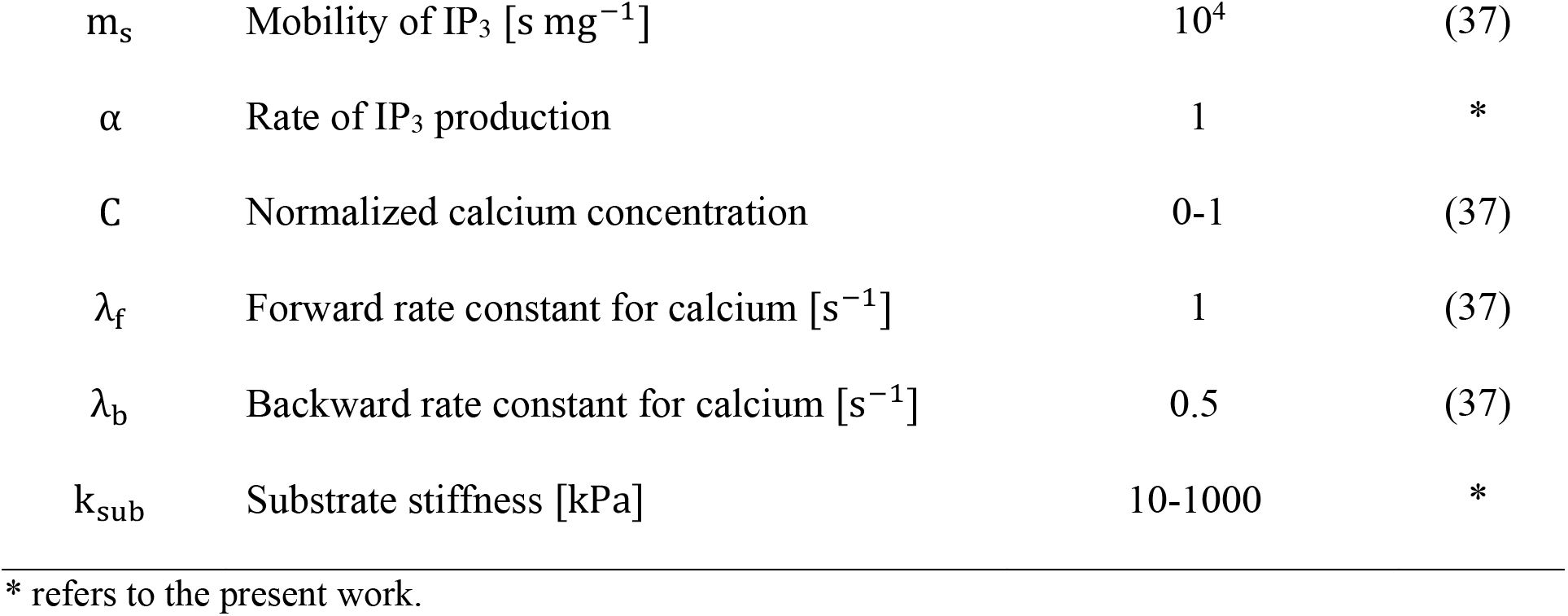
List of variables and values used in the stochastic FE composite model

#### 6. Model simulations and parametric analysis

We quantified the effects of ligand density and substrate stiffness on cell-substrate interactions under static and stretch conditions. Varying ligand densities corresponding to 420, 740, 1060, and 1380 μm^-2^ were used and the model simulated at fifteen different substrate stiffness values ranging between 10-1000 kPa to quantify cellular responses. The static (no-stretch) case was used as a control for each of these simulations by imposing zero displacements at the ends of the substrate. These simulations were compared with the cyclic stretch condition of 10% at 1 Hz applied to the substrate. Cell-substrate interactions were evaluated for ~10^5^ iterations in the stochastic simulations with each condition repeated twenty times. Model outputs include the computed cell tractions at each location of the cell, integrin dynamics, calcium concentrations, and clutch engagements. Output values were averaged spatially and temporally to explore the combined effects of substrate stiffness and stretch on SF and FA dynamics.

## Results and Discussion

Quantification of cell-substrate interactions under cyclic stretch involves coupling the SF contractility and activation, dynamics of clutch interactions (adaptor proteins) with the substrate, and calcium signaling. We combined these individual modules into a chemomechanical model of the fibroblast in our study using a novel SFEM framework (Fig. 2; Table 1). We investigated the individual and combined roles of SF contractility and calcium signaling on clutch engagements/ dis-engagements. We also quantified the effects of varied substrate stiffness and cyclic stretch on cell adhesions and tractions using the model.

There are three main results from this study: first, we show that integrin recruitments changed along the length of the cell and were low at lamellipodial regions located at the leading edge of the cell. The highest adhesions were in the lamellar region located some distance away from the leading cell edge. Intracellular calcium resulting from FA dynamics was highest at the lamellipodium. Second, cyclic stretch altered the SF contractility, calcium, and tractions as compared to the static (no-stretch) case. The overall SF contractility increased and was accompanied with a corresponding decrease in the integrin recruitments for stretched cells. Finally, cell tractions and adhesions show biphasic responses with substrate stiffness and increased with higher substrate ligand density.

### Effects of SF dynamics and calcium signaling on motor-clutch dynamics

We simulated the effects of calcium signaling and SF contractility on traction generation at focal adhesions for a fibroblast on 10 kPa substrate coated uniformly with 420 μm^-2^ fibronectin density.

We explored three different cases at the lamellipodia to assess the influence of varying SF activation and calcium signaling in the model. (i) The motor-clutch model with constant SF activation (η=1) and calcium (C=1). This case served as a control to assess the influence of differential SF activation and calcium signaling in the model. (ii) The motor-clutch model with exponentially decaying calcium (time constant, θ = 720 s) illustrates changes in SF remodeling with calcium. (iii) Motor-clutch model with calcium feedback due to integrin engagements/ clustering corresponding to SF remodeling caused by the resisting forces of clutches.

Figure 3A shows temporal variations in the SF activation due to calcium signaling that leads to changes in the motor-clutch dynamics. SF activation is lower than the control case when the SF is always activated and maintained at a constant value. We quantified the retrograde flow velocity of actin (*v*), clutch engagements (*n_eng_*), and substrate traction *(T)* for all three cases. Clutches undergo load-fail behaviors and vary cyclically between extrema. These variations are a function of SF contractility and the load-bearing capacity of the clutches (Fig. 3B, C, and D). An increase in the force on the *i*^1h^ clutch, (*F*_c[*i*]_), results in an increase in the clutch off-rate according to the unloaded off-rate (*k_off_* = 0.007 s^-1^) and characteristic rupture force, *F_b_* (Eq. 1). The retrograde actin flow varied between 10 – 120 nm/s. Tractions in the motor-clutch model varied between 0-12 nN (Fig. 3B, D). Experiments show a similar range of fibroblast traction with average traction of 3 nN/ μm^2^ on a continuous substrate (5,11). The range of forces applied at an adhesion site varied between 0-10 nN. Because the SF activation is a constant in the control case (Fig. 3A), the magnitude and frequency variations (~0.018 Hz) in load-and-fail of clutches remain essentially constant. The retrograde actin flow and tractions are inversely related such that the traction maxima corresponds to the minima for retrograde flows. Forces build in the clutches and fail catastrophically (Fig. 3C) when the ensemble reaches load-bearing capacity.

**Figure 3:**
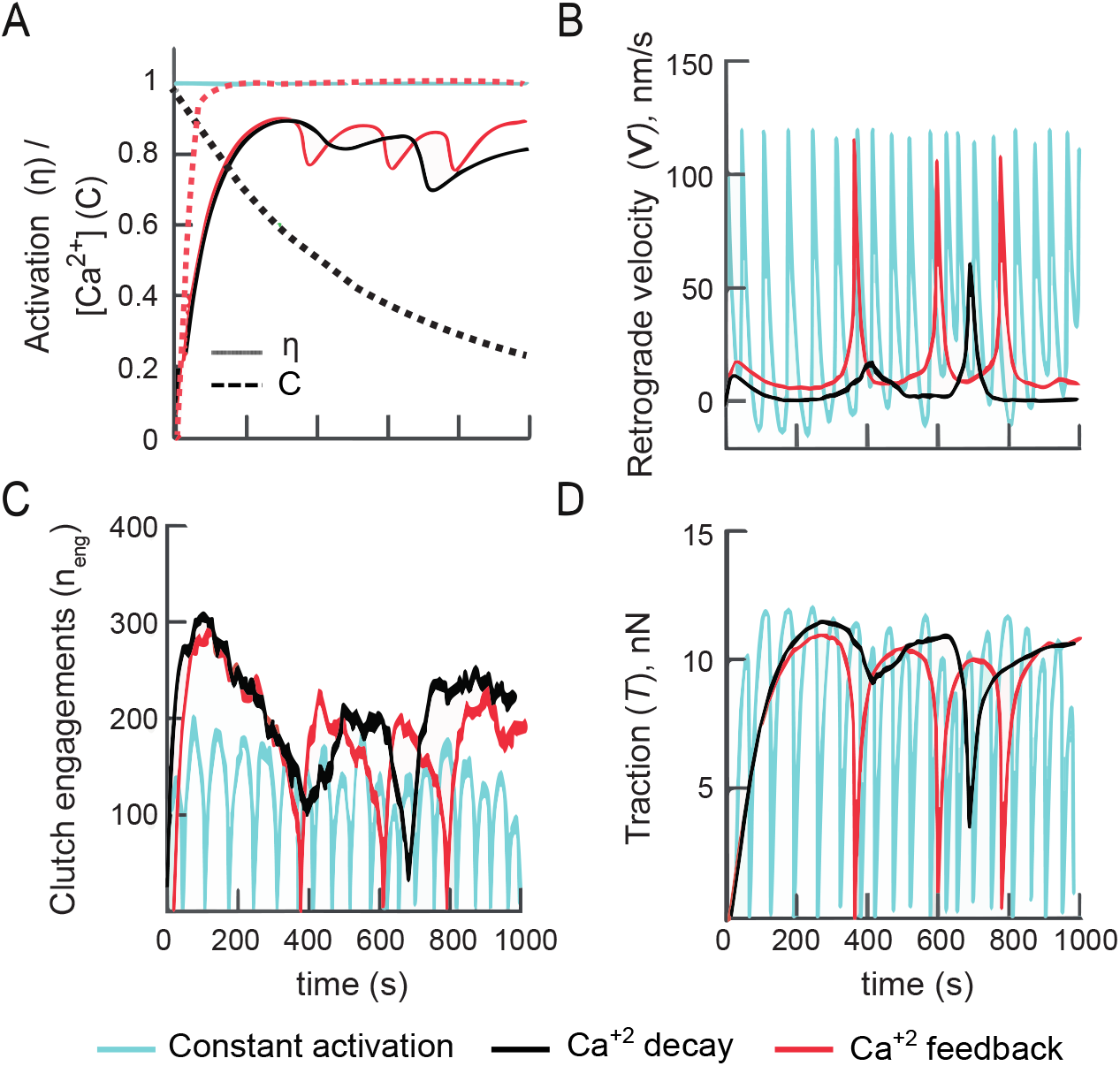
Motor-clutch dynamics are shown at the lamellipodia corresponding to (i) constant activation (η, C =1), (ii) exponential calcium decay, and (iii) calcium feedback due to integrin engagements/ clustering and SF remodeling. **(A)** Temporal evolutions in the SF activation (solid) and calcium are shown for the three different cases (dotted), **(B)** Retrograde actin flow velocity shows a load-and-fail profile. The variational frequency is dependent on the temporal evolution of SF activation and calcium feedback. **(C)** Clutch engagements are higher and show lower load-and-fail events with decrease in calcium concentration. **(D)** Tractions exerted on the substrate bear similarities with the profile for retrograde flow. Maximum tractions were obtained for minimum values of the retrograde flow.

In the second case, SF activation decreased with decaying calcium over time due to decreased calcium (Eq. 6) and cyclical clutch failures (Fig. 3A, C). Retrograde actin flow (*v*) and tractions *(T)* reached maximum values of 12.5 nm/s and 11.75 nN respectively (Fig. 3B, D). Cyclic variations, clearly identified in the control case, are not as apparent because the SF is not maintained at a maximum value of activation. Decreased SF activation resulted in higher clutch engagements and a corresponding decrease in actin retrograde flow. The velocity does not hence reach the free control velocity value of ~120 nm/s. The model reaches a stalled regime, characterized by near-zero retrograde actin flow, which decreases the cytosolic calcium. The number of engaged clutches was higher with lower retrograde actin flow (Fig. 3A, C).

The influx of calcium from the ER in the third case maintains calcium feedback in the cell (Eqs. 8, 9, and 11). Cyclic variations in clutch engagements resulted in changes to the concentrations of high-affinity integrins (Eq. 9) and were involved with the release of calcium. SF activation, clutch engagements, and the associated tractions varied such that the system did not reach a stalled state (Fig. 3A, C). Fluctuations in the SF activation are a result of variations in the reaction forces of clutches. These simulations show that the calcium signal and SF activity play an important role in the magnitude of cell tractions (Fig. 3D) and retrograde actin flows at the FA regions.

The dynamics of adaptor protein engagements at the FA sites in the motor-clutch model vary with SF contractile force (25). Clutches, represented as slip bonds, reversibly engage with actin through myosin dependent processes. Three separate regimes characterize these interactions: first, *frictional slippage* at low stiffness with an intermediate magnitude of retrograde flow and traction forces. Second, the *load-and-fail* regime at intermediate stiffness with a slow retrograde flow and higher traction forces, and finally, *frictional slippage* at high stiffness with high retrograde flow and low tractions (26). Lower resistance to actin flow was observed in the frictional slippage region on compliant substrates due to clutch failures before the SF could achieve a maximum possible contractile force. Clutches prematurely fail on compliant substrates as deformations occur at a slower pace than the SF contraction. In contrast, stiff substrates have rapid clutch deformations that result in failures due to lower substrate deformations. The load-and-fail regime was associated with clutches undergoing maximum deformations that collectively fail with SF contraction (Fig 3C). We see a similar delineation of the different regimes in our study. We emphasize that these regimes were simulated on a single substrate with the incorporation of SF remodeling and calcium signaling that has not been discussed in earlier motor-clutch models. Our results suggest that modulations in the cytosolic calcium helps tune cellular tractions at FA’s.

### Spatiotemporal variations in cell-substrate interactions with cyclic stretch

Cyclic stretching caused dynamic variations in the cellular responses. We quantified changes in actin retrograde flow (*ν*), clutch engagements (*n_eng_*), traction (*T*) and calcium signaling at different spatial locations within the adherent cell on a substrate coated with a uniform fibronectin ligand distribution. The substrate had a uniform ligand density, n_c_= 420 per μm2, and has a modulus, *E_sub_* = 10 kPa. Simulations were compared for static and cyclic sinusoidal stretch (10% amplitude with 1Hz frequency) for 10^5^ events. The ends of the substrate were initially fixed and then stretched during the subsequent half of the simulation.

Figure 4 shows the time profile of cell-substrate interactions at the cell interior located ~7.24 μm from the leading edge. Actin retrograde flow velocity and tractions show load-and-fail characteristics under no-stretch conditions with variations between 0-60 nm/s and 0-10 nN at ~5 Hz frequency (Fig. 4A and D). These results are similar to the motor-clutch simulations with calcium feedback and are associated with a change in chemical signaling represented by IP_3_ and calcium concentrations (Fig. 4C).

**Figure 4.**
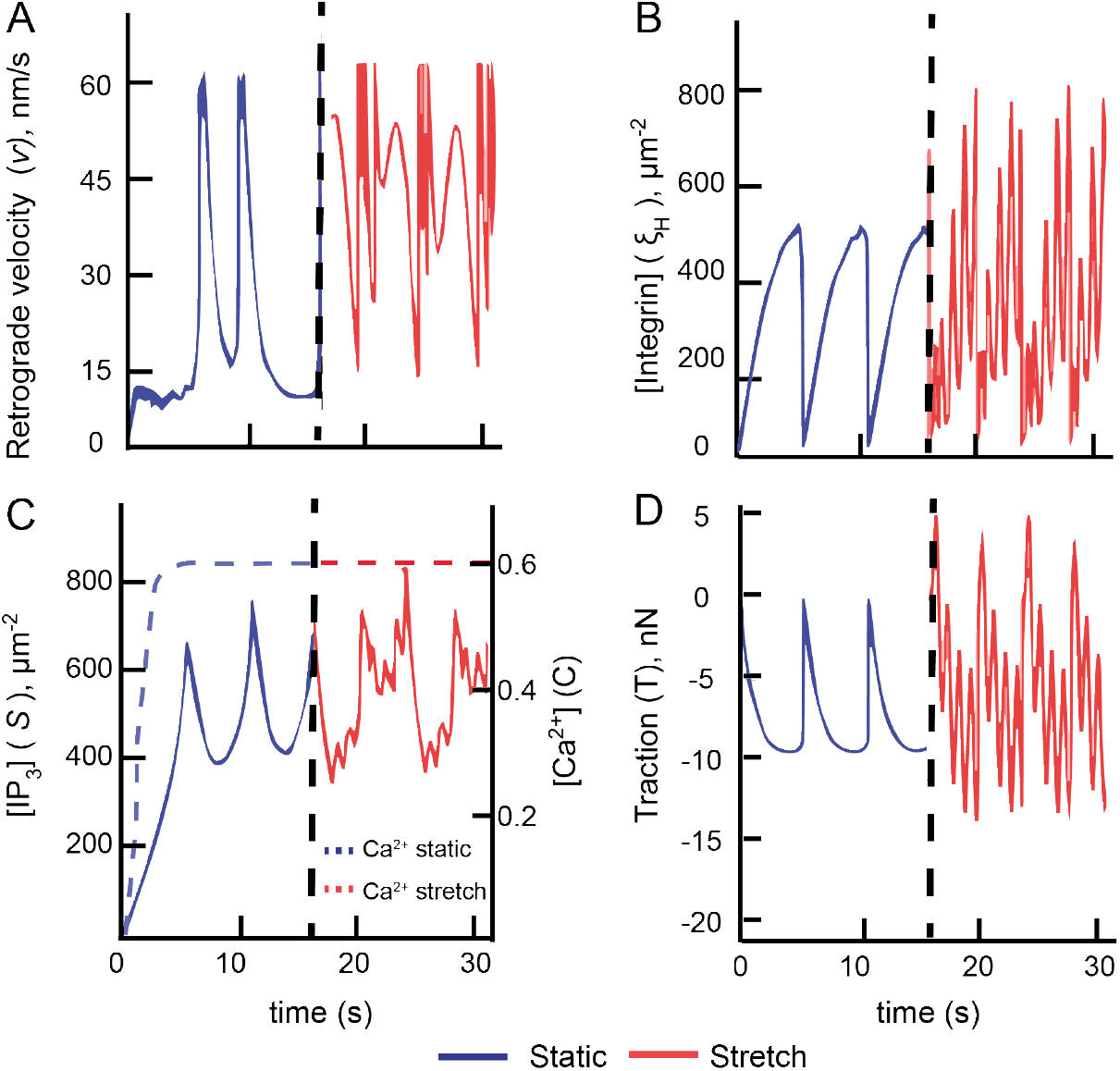
Temporal evolutions in cell adhesions under static and 10% stretch are shown at node 7 located ~7.24 μm from the leading cell edge. 10^5^ events were simulated on a 10 kPa substrate with 420 μm^-2^ fibronectin concentration. **(A)** The retrograde actin velocity for the static case increases with time due to SF activity. Retrograde flows are higher under stretch due to higher SF activation induced by the high-affinity integrin recruitment. **(B)** The concentration of high-affinity integrins follows load-and-fail behavior with higher variations under stretch. **(C)** IP_3_ production increased with integrin engagement density. Calcium increased with IP_3_ concentration and plateaued at a constant level under stretch. **(D)** Changes in cell tractions resemble the integrin engagement profiles and have maximum values at minimum values of the retrograde velocity.

Initial clutch engagements cause an accumulation of high-affinity integrins (Fig. 4B) which results in the release of IP_3_ and calcium. Clutch engagements resist the retrograde flow as forces build in the clutches. Excessive deformations cause clutch failures that result in cyclic variations in the clutch forces (*F*_c[*i*]_) and corresponding changes in the concentrations of high-affinity integrins (*ξ*_*H*[*i*]_) (Fig. 4B). Variations in the rate of IP_3_ production due to a change in the integrin concentrations (Eqs. 9 and 10) result in IP_3_ concentrations changes between two extrema. Calcium released by IP_3_ increased monotonically and plateaued at later time durations.

Substrate stretch induces additional forces in clutches that cause perturbations in tractions and high-affinity integrin concentrations. Higher forces in the engaged clutches resulted in higher SF activation and clutch disengagements (Eq. 1) that were accompanied by higher average actin retrograde flow velocity (Fig. 4A). The increase in the integrin density under cyclic stretch resulted in higher production of IP_3_ and a corresponding increase in the cytosolic calcium (Fig 4B, C). Calcium increase, deformation-induced higher tractions in the cell, and the inverse correlations between traction and retrograde, were also observed under cyclic stretch conditions similar to the control (no-stretch) case. The number of load-and-fail events were however higher under stretch (Fig 4A and D). Additional clutch deformations under stretch resulted in ~50% increase in the traction magnitude and ~ 33.3% increase in high-affinity integrin concentrations.

Figure 5 shows variations in tractions, clutch engagements, high-affinity integrins, and calcium concentrations along the cell length for static and 10% cyclic stretch cases. The asymmetric traction profile is a consequence of the sign convention employed in the study. Tractions were higher in the transition region adjacent to lamellipodial regions. Lower tractions at the cell center were due to the lower recruitment of high-affinity integrins and greater clutch attachment failures in the lamellipodia. Calcium was higher at the cell center and edges. Tractions, high-affinity integrins, and calcium concentrations have similar profiles under cyclic stretch as the static case but with significantly higher values.

**Figure 5.**
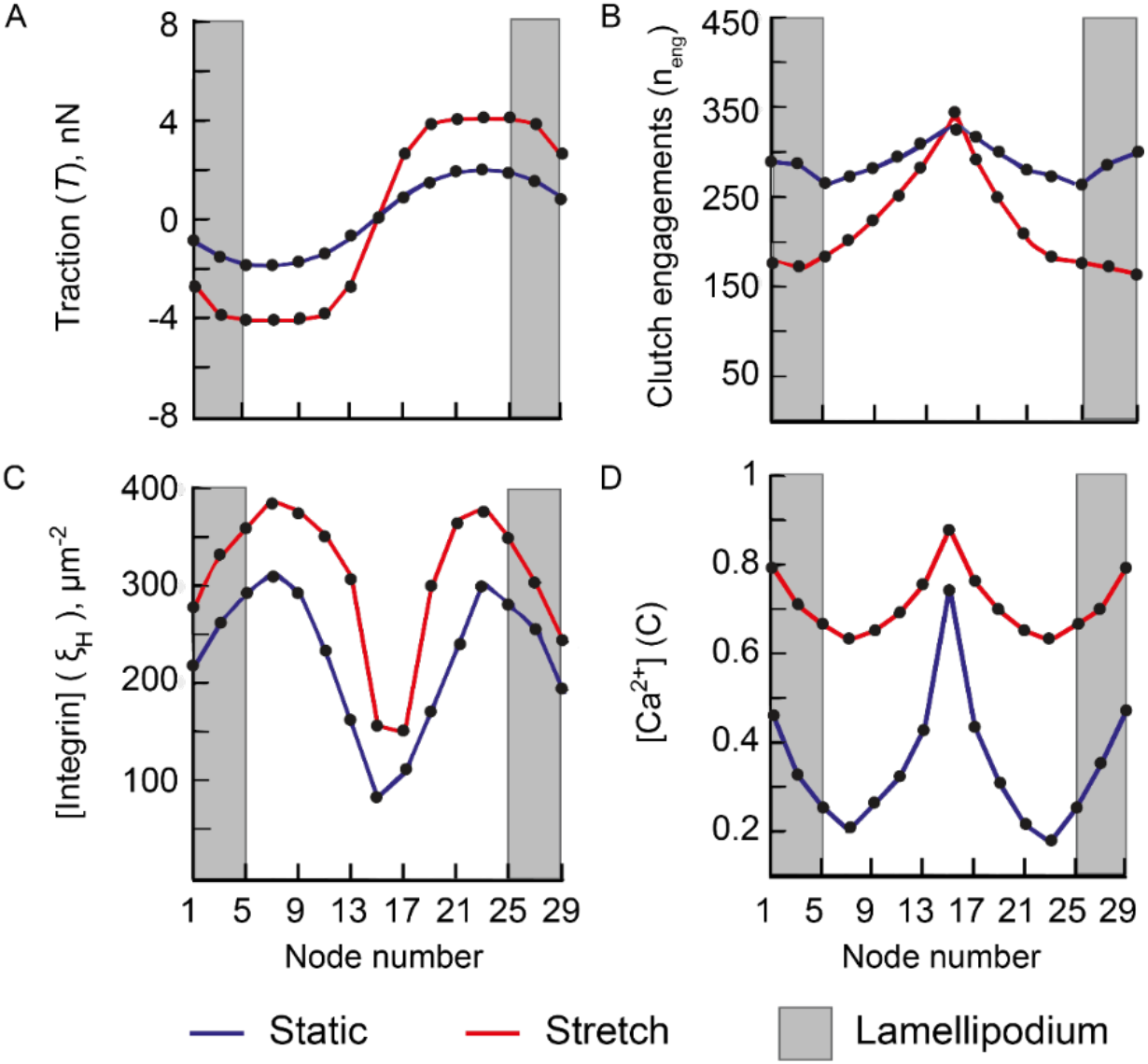
Spatial variations in **(A)** tractions, **(B)** clutch engagements, **(C)** high-affinity integrin concentration, and **(D)** calcium concentration over the cell length for static and cyclic stretch conditions.

Interactions between the SF and reversibly engaging clutches along the cell resulted in SF and FA’s remodeling and a corresponding release in calcium signaling molecules. These results show the emergence of three distinct regions in the cell: the lamellipodial regions in gray, the transition region, and the central region of the cell.

The traction profile was antisymmetric about the node 15 corresponding to the cell center (Fig 5A). Tractions result from the combined effects of SF contractility and the resistance provided by engaged clutches to actin retrograde flow. A maximum traction of ~1.93 nN was present at nodes 7 and 23 of the cell in the transition region for the static case (Fig. 5A). Lower tractions were observed at the lamellipodial regions (1-5 nodes) whereas the cell center had near-zero tractions. The retrograde flow of actin, induced by the contractile strain in the SF, decreased monotonically from the lamellipodial region to the center due to the resistance of clutch engagements. A lower retrograde velocity induces optimum deformation of clutches at lamellar regions which resulted in higher tractions (Fig 5A).

Experiments report decreased actin retrograde flow velocity from the lamellipodium region to the lamellar regions of the cell (41). Embryonic fibroblasts show an average actin retrograde flow of 11.33 nm/s in the lamellipodial regions and 5.5 nm/s in the lamellar regions. Tractions varied cyclically between ~2.5-1.89 nN at the FA. Because a majority of the load in the SF was transferred to clutches in lamellar regions that develop maximum tractions, the resulting force transfer at the cell center was lower as compared to lamellipodial regions. Similar spatial variations in tractions were reported in fibroblasts seeded on polyacrylamide substrates; the maximum traction was 4 nN/μm^2^ at a distance ~6 μm from cell edge (42). In contrast, fibroblasts on discontinuous pillar-ridge substrate have the highest force (~14 nN) at the cell edges (43). Epithelial ptK1 cells also show spatial changes in the actin retrograde velocity and tractions (44). Retrograde velocity monotonically decreased from the cell edge (~15 nm/s) to the center (~1.5 nm/s). Tractions increased from ~25 kPa at the cell edge to ~90 kPa in the intermediate region and were near zero towards the cell center.

Simulations show that cyclic stretching of the substrate have a similar traction profile as the static case. Tractions under stretch increased by ~50% as compared to the static case (Fig 5A). The overall tractions increased to ~4 nN in the intermediate region and ~2.76 nN at the cell edge. Higher tractions were a result of higher clutch deformations and calcium induced by external stretch to the substrate. Integrin concentrations increased under cyclic stretch; values were lower in the lamellipodial regions and increased towards the lamellar regions (Fig. 5B). High clutch engagements at the cell center are due to a lower transfer of force through clutches (Fig 5A). A combination of low tractions and high clutch engagements leads to a stall at the cell center. The lamellipodial and intermediate regions have high tractions and clutch deformations with load-fail profiles (Fig. 4). Cell-substrate interactions varied from the cell edge to the center. These results are contrary to those reported earlier by Deshpande and colleagues using a model for SF activation and integrin remodeling (37). The study showed higher integrin density and tractions at the cell edge that are contrary to experimental studies and the results from our simulations. These differences may be due to the non-inclusion of slip bond characteristics in the integrin-fibronectin bond that may lead to an increase in the force transfer between cell and substrate at the lamellipodial regions.

The higher accumulation of high-affinity integrins (*ξ_H_*) at the lamellar and lamellipodium region correlated with higher traction magnitudes (Fig. 5C). Integrin accumulation at the cell center is primarily driven by a difference in the reference chemical potential between high-affinity and low-affinity integrins due to lower tractions. Calcium concentrations show a symmetric profile about the cell center (Fig. 5D) with higher values at the lamellipodial regions and cell center as compared to the static case. Calcium was higher at lamellipodial regions due to the frequent failure of clutches that cause higher IP_3_ (Fig. 5B). Cyclic stretch caused an increase in the intracellular calcium with similar variations along the cell length as compared to the static case; these results agree with experiments on cyclically stretched fibroblasts (45).

### Cellular mechanosensitivity to substrate stiffness

We used the model to investigate the role of substrate stiffness and ligand density on cell adhesions under static and cyclic stretch. Fig. 6 shows variations in the magnitude of average substrate tractions, clutch engagements, high-affinity integrin, and calcium concentration computed over the entire cell length. We simulated various ligand densities (420, 740, 1060 and 1380 *μm*^-2^) and used fifteen distinct substrate stiffness (10-1000 kPa) to represent a range of *in vivo* tissue stiffness (46). The mean clutch forces and high-affinity integrin concentrations were used to compute the average tractions, *T_sub_,* and the average integrin concentrations in the study.

**Figure 6:**
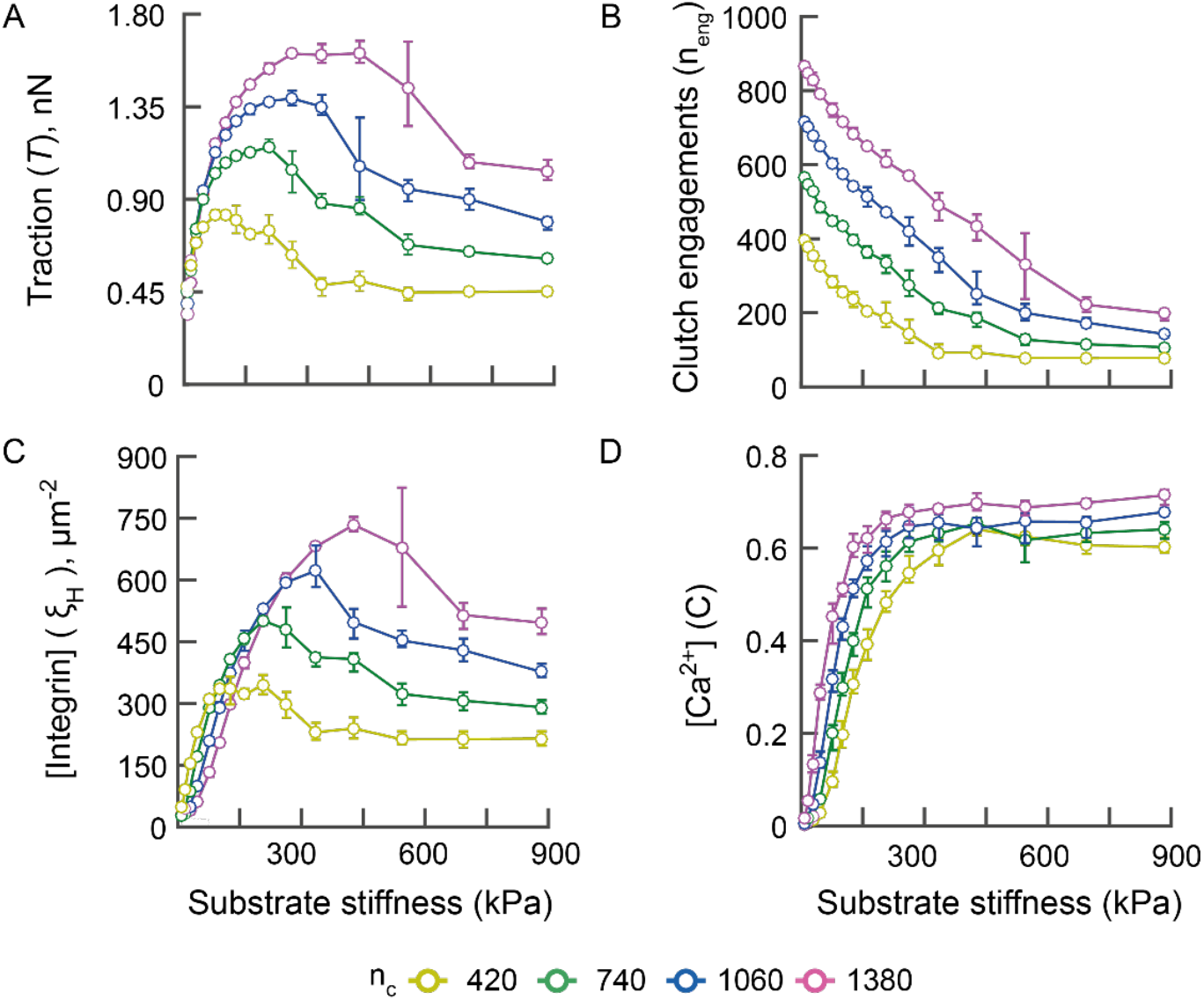
**(A)** Cell tractions peak with substrate stiffness under static (no-stretch) condition for each ligand density and plateau to a constant value at higher stiffness for each of the different ligand densities in the study. **(B)** Clutch engagements, n_eng_, decreased with substrate stiffness. Higher engagements were visible on substrates coated with greater ligand density. **(C)** Integrin concentrations, *ξ_H_*, shows a biphasic response with substrate stiffness. **(D)** Calcium concentrations increased monotonically with substrate stiffness and saturated for each ligand density.

Data were averaged from 20 simulations at each substrate stiffness and ligand density. Average values, along with variations, are shown using error bars in the figure. Cell tractions had a biphasic relationship with the substrate stiffness, *k_sub_* (Fig. 6A). Tractions increased initially with substrate stiffness, reached a maximum value, and subsequently decreased. High clutch engagements occur between the SF and the substrate at very low values of stiffness and corresponding to each ligand density (Fig. 6B). The presence of additional ligands at high concentrations permit more clutches to bind which increases the load-bearing capability of individual clutches and results in higher tractions (Fig. 6A, B). The (average) high-affinity integrins also display a biphasic relationship with substrate stiffness similar to that of cell tractions (Fig. 6C). An increase in the ligand density on the substrate permits greater integrin accumulations sites and higher integrin recruitments. The associated increase in clutch forces decreases the chemical potential of high-affinity integrins and result in higher tractions (Eq. 9). Substrates with stiffness less than 3 kPa, and with low ligand densities, have higher integrin concentrations.

These results put into focus the effects of clutch forces on integrin densities that may outweigh the presence of additional integrin accumulation sites on substrates with low stiffness (Fig. 6C). Ligand density and substrate stiffness play an important role in FA formation in fibroblasts and myoepithelial cells (47, 48).

Higher ligand density and substrate stiffness monotonically enhanced cell spreading and integrin recruitments at FA in fibroblasts. Larger FA regions were observed in myoepithelial cells at lower ligand densities on substrates with stiffness < 30 kPa; FA’s are destabilized on stiffer substrates. These experiments demonstrate the biphasic nature of integrin recruitments with substrate stiffness. Our results also demonstrate the sensitivity of calcium concentration to substrate properties. Fig. 6D shows a rapid increase in calcium at lower substrate stiffness which saturates to a maximum value for each of the different ligand densities in the study. The calcium concentrations remain relatively constant on stiffer substrates. Cell mechanosensitivity to substrate stiffness may hence be tuned by altering the ligand densities on the substrate that alters the calcium signaling in cells.

### Cyclic stretching increases cellular tractions and integrin recruitments at FA

We investigated cyclic stretch effects on cell tractions, adhesions, and calcium concentrations on substrates with varying ligand densities and stiffness using the composite model. The (average) substrate tractions, *T_sub_*, and high affinity integrin concentrations have a biphasic relationship with substrate stiffness that is similar to no-stretch conditions. These variations have a clearly defined and a relatively sharp peak as compared to the corresponding values in the static case (Fig. 7). Higher ligand densities resulted in increased tractions, clutch engagements, high-affinity integrins, and calcium in cells. Although traction magnitudes under cyclic stretch were significantly higher on compliant substrates as compared to the static case, the average tractions were lower on the stiffer substrates and reduced to near zero values. The corresponding values of clutch engagements and high-affinity integrin densities were also significantly lower under cyclic stretch. The maximum cell traction under cyclic stretch (n_c_ =1380 *μm*^-2^) was ~50% higher than the corresponding static case; ntegrin concentrations were also 63% higher under stretch as compared to the static case.

**Figure 7.**
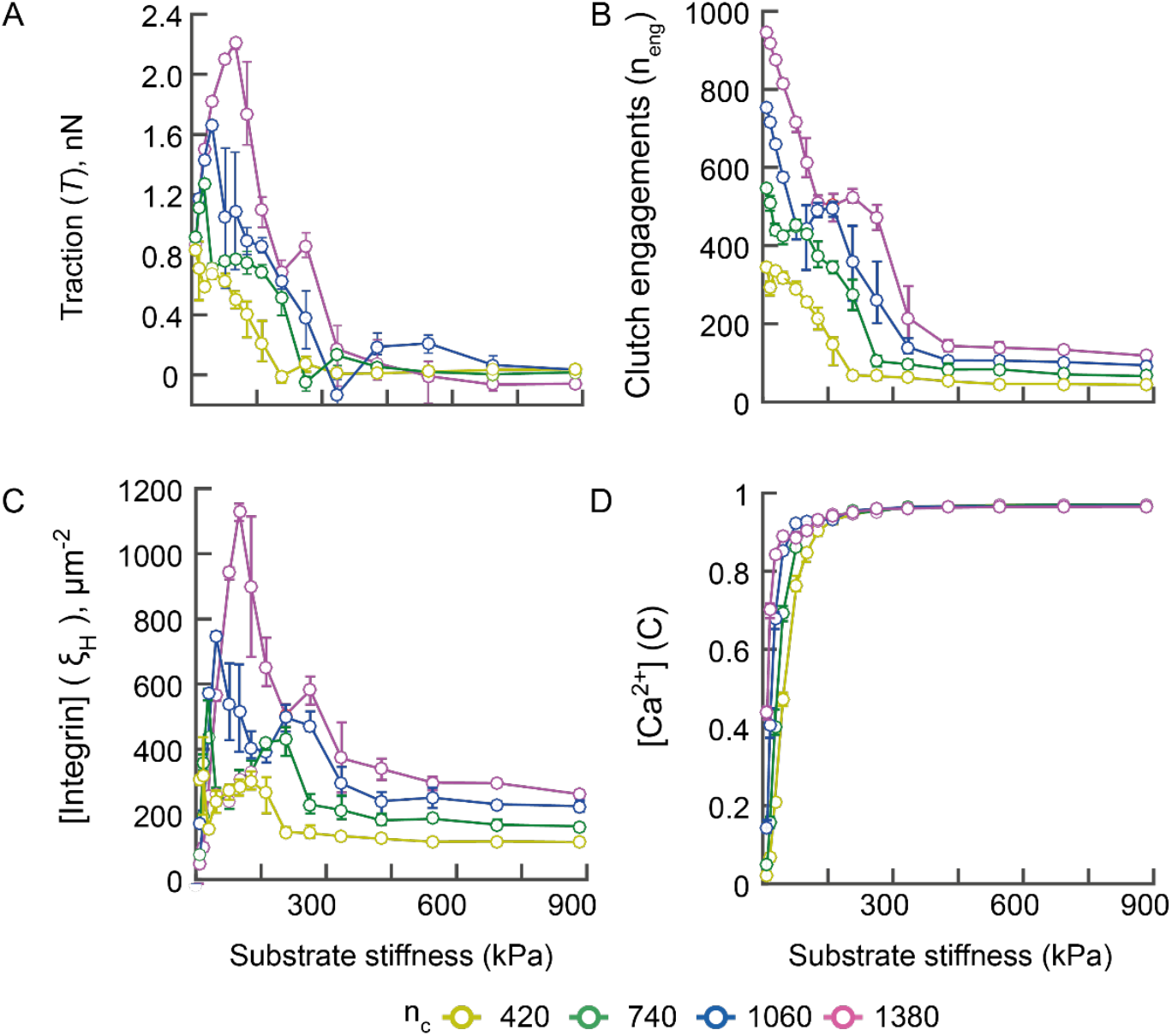
**(A)** Variations in the average cell tractions with substrate stiffness under cyclic stretch consitions. Tractions reaches maximum values at an optimal substrate stiffness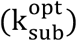. **(B)** Clutch engagements decreased monotonically with substrate stiffness for each of the different ligand densities. **(C)** Integrin concentrations for engaed clutches, *ξ_H_*, show biphasic response with substrate stiffness. **(D)** The calcium concentration increased monotonically with substrate stiffness and reached a saturation value for each ligand density. Variation in ligand densities do not have a significant effect on cytosolic calcium. Data were averaged from 20 simulations at each substrate stiffness and ligand density. Average values, along with variations, are shown using error bars in the figure.

Clutch engagements monotonically decreased with substrate stiffness at a fixed ligand density as seen earlier for cells under no-stretch condition. These values were lower under stretch due to the higher tractions. (Fig. 7B).

An increase in ligand density shifts integrin recruitments to higher values; integrin concentration in the cell approaches near-zero on substrates with stiffness beyond the critical stiffness (Fig 7C). Calcium concentrations also increased monotonically with substrate stiffness for all ligand densities similar to the static case, and reached a maximum level after ~200 kPa under cyclic stretch (Fig. 7D).

Experiments demonstrate that fibroblasts subjected to 5% cyclic stretch at 1 Hz frequency have 66% increased tractions (49). A higher SF activation, caused by clutch deformations induced by stretch, may be a primary reason for the underlying difference between the static and cyclic cases. Substrate stretching resulted in an increase in calcium concentration due to the higher *IP*_3_ concentrations caused by changes to the high-affinity integrin density, *ξ_H_* (Fig. 7C). Application of 7% stretch on vascular smooth muscle cells increased calcium concentration from ~85 nM under no-stretch to ~95 nM under cyclic stretch (52). Model predictions from our study demonstrate that increased tractions are mechanistically linked to higher SF activity and coupled to increased calcium and the recruitment of high-affinity integrins at FA in cells under cyclic stretch.

### Optimal substrate stiffness sensing of fibroblasts to cyclic substrate stretch

We quantified substrate stiffness sensing of fibroblasts using cell tractions for cyclic stretching of substrates with varying ligand densities. An optimal substrate stiffness 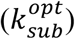 was defined based on the maximum (average) traction at a given ligand density. Results for 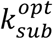 were compared between static and cyclic stretch cases for each of the different ligand densities in the study. Simulations from the composite model show that the maximum traction at 420 μm^-2^ ligand density was similar for static and stretch conditions and increased by ~40% for 1380 μm^-2^ ligand density (Fig. 8A).

**Figure 8.**
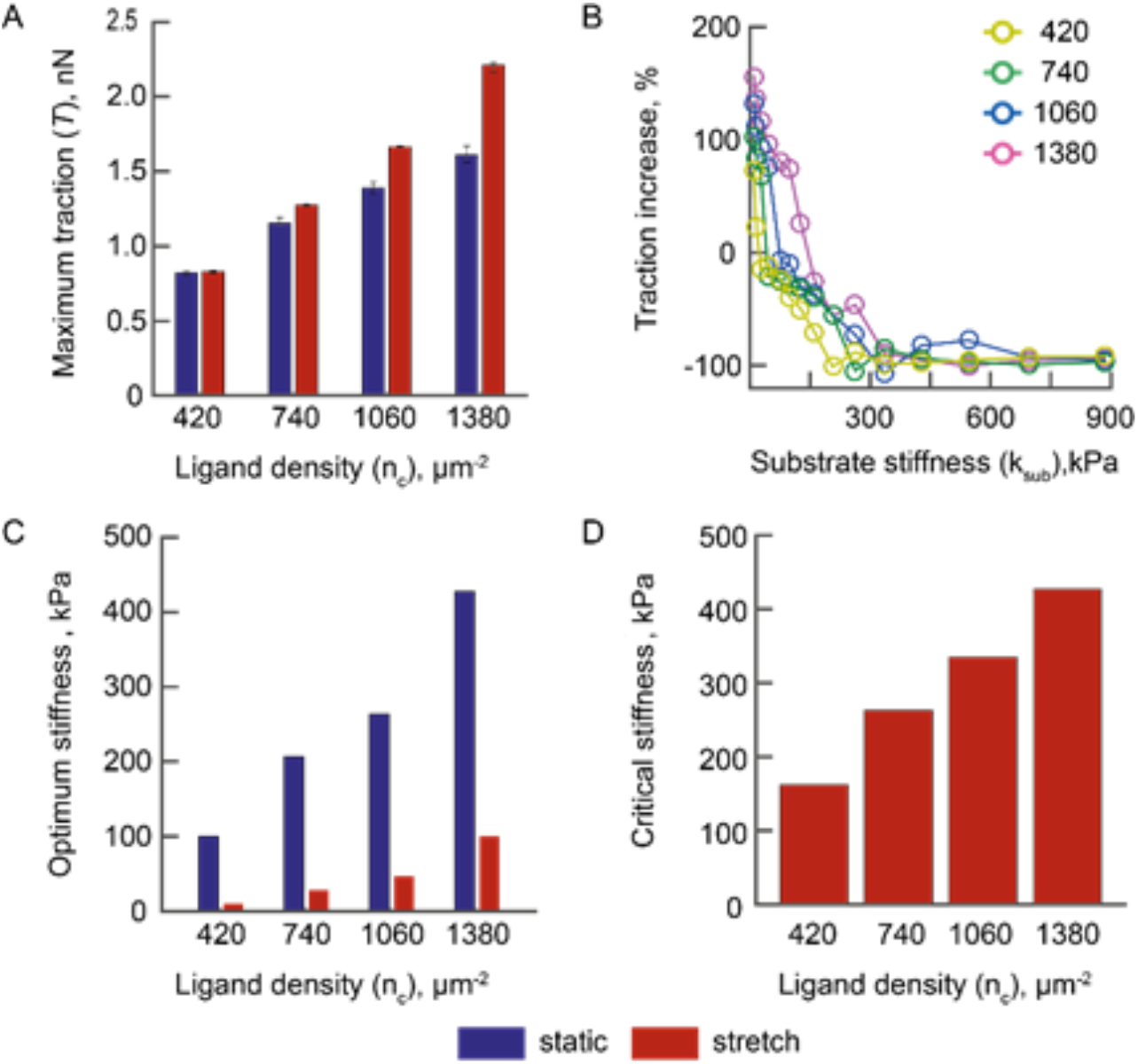
(**A**) Maximum traction, *T*, variations as a function of ligand density are shown for 10% cyclic stretch and static conditions. (**B**) The increase in the traction percent due to stretch as a function of substrate stiffness is plotted for different ligand densities. Substrate stretching increases cell tractions on compliant substrates; the cell tractions however decrease on stiff substrates. (**C**) Optimum stiffness, computed corresponding to the maximum tractions, are plotted as a function of ligand density. An increase in ligand density shifts the optimum stiffness towards stiffer values in both static and stretch cases. (**D**) Critical stiffness, characterized by near-zero tractions, increases with ligand density due to stabilization of the adhesions.

We quantified percent increase in average traction due to application of stretch as

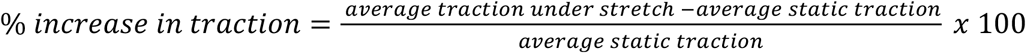

Figure 8B shows enhanced tractions under cyclic stretch for all ligand densities corresponding to compliant substrates in the study; tractions were significantly lower on stiff substrates. An increase in the ligand density resulted in higher traction magnitudes over a larger substrate stiffness range. Premature failures of adaptor proteins, connecting the SF and the FA, at low substrate stiffness were prevented under stretch which results in higher tractions. Stretching of substrates with higher stiffness caused clutch overloading which shifts cell-substrate interactions into the frictional-slippage regime and results in lower tractions.

The optimum stiffness shifted towards compliant substrates for all the ligand densities in the study for cells under cyclic stretch (Fig. 8C). The magnitude of shift was greater for higher ligand densities. For example, the optimum substrate stiffness shifted by 90 kPa at 420 *μm*^-2^ and by 300 kPa for the 1380 *μm*^-2^ ligand density. Stretching of substrates stiffer than the critical stiffness significantly reduces fibroblast tractions and integrin recruitments at ligands (Fig. 7A and 7C). The availability of additional ligands stabilizes adhesions to the substrate and results in an increase in the critical substrate stiffness (Fig. 8D). Near-zero values of tractions also demonstrate cell de-adhesion under stretch at various ligand densities.

Chan *et al.* used the motor-clutch model to show an optimal stiffness during cell-substrate interaction at FA (26). Experiments demonstrate the biphasic relation between average tractions and substrate stiffness for fibroblasts (53). They observed maximum traction of 1.5 nN on ~15 kPa substrates. Results from our study show a similar biphasic tractions with a maximum value of 1.35 nN at 1060 *μm*^-2^.

We considered only one type of integrin (α_5_β_1_) in the model and did not include the effects of force loading rate on integrin-fibronectin bond. We have also not included the effects of actin polymerization on adaptor protein assembly. Coupling of chemical signaling with mechanical properties of cell adhesions to substrates clearly demonstrates the effects of SF and FA remodeling under cyclic stretch that has not been shown earlier. A quantification of fibroblast adhesions under cyclic stretch is essential to our understanding of mechanobiological changes in diseases like arterial fibrosis.

## Conclusions

We use a systems biology approach to develop a one-dimensional, multi-scale, stochastic finite element model of cellular adhesions to linear elastic substrates of varying stiffness and different ligand densities. The model includes FA attachment dynamics, SF activation, and calcium signaling. We quantified the resulting cellular tractions in lamellipodial and other cellular regions in response to variations in substrate stiffness, cyclic stretching, and ligand densities. Calcium signaling is essential in modeling cell – substrate interactions. A high concentration of calcium enhances SF activation and permits integrin assembly at FA regions. Our model shows that mechanical interactions between the SF and reversibly engaging adaptor proteins at FA contribute to the spatiotemporal variations in cell contractility and adhesions reported in experiments. Tractions and integrin recruitments vary along cell length and have maximum values in the lamellar regions of the cell. The optimal substrate stiffness at which cells exert maximum tractions, shifted towards stiffer substrates at high ligand densities. Cytosolic calcium also increased with substrate stiffness and ligand density. We explored changes in the cell behaviors under 10% cyclic stretch at 1 Hz and compared these responses with static (no-stretch) condition. Cyclic stretch resulted in increased cytosolic calcium and corresponding higher tractions at the lamellipodial and intermediate regions of the cell. Cell tractions and adhesions have a biphasic variation with substrate stiffness and increase with higher ligand densities. The optimal substrate stiffness at a given ligand density shifted towards compliant substrates with cyclic stretch. Integrin concentrations and tractions reduce to near-zero values beyond the critical substrate stiffness; these results show cell deadhesion under cyclic stretch. Chemomechanical coupling is essential in the mechanosensing responses underlying cell-substrate interactions.

## Author Contributions

SJ performed the simulations, analysed the results, and helped write the manuscript. NG designed the study, supervised the research, helped with data analysis, and wrote the manuscript.

## Acknowledgements

NG gratefully acknowledges the Department of Science and Technology (SERB/003640) for project support.

## Supplemental Information

**List of Supplementary Figures**

**Figure S1:**
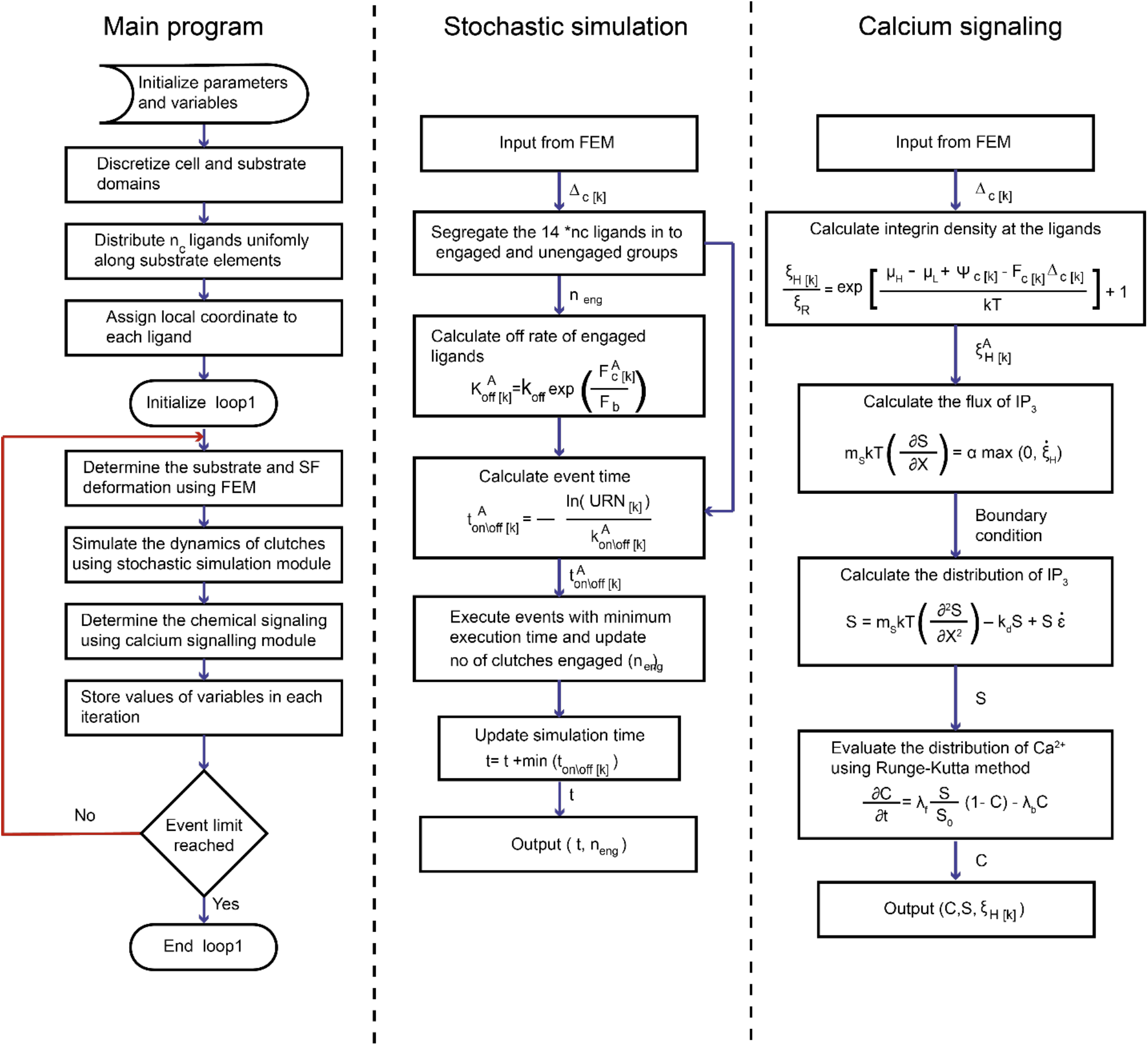
The flow chart aims to outline the sequence of events in the simulation clearly. The main program evaluates the mechanical equilibrium, stochastic simulation, and calcium signaling modules during cell-substrate interaction during each time step. The events in these modules are shown separately, with corresponding inputs and outputs on the arrows.

